# Disentangling environmental effects in microbial association networks

**DOI:** 10.1101/2021.07.13.452182

**Authors:** Ina Maria Deutschmann, Gipsi Lima-Mendez, Anders K. Krabberød, Jeroen Raes, Sergio M. Vallina, Karoline Faust, Ramiro Logares

**Author notes:** Corresponding authors: Ina Maria Deutschmann, Karoline Faust, Ramiro Logares. Shared last authors.

## Abstract

**Background:** Ecological interactions among microorganisms are fundamental for ecosystem function, yet they are mostly unknown or poorly understood. High-throughput-omics can indicate microbial interactions through associations across time and space, which can be represented as association networks. Associations could result from either ecological interactions between microorganisms, or from environmental selection, where the associations are environmentally-driven. Therefore, before downstream analysis and interpretation, we need to distinguish the nature of the association, particularly if it is due to environmental selection or not.

**Results:** We present EnDED (**En**vironmentally-**D**riven **E**dge **D**etection), an implementation of four approaches as well as their combination to predict which links between microorganisms in an association network are environmentally-driven. The four approaches are Sign Pattern, Overlap, Interaction Information, and Data Processing Inequality. We tested EnDED on networks from simulated data of 50 microorganisms. The networks contained on average 50 nodes and 1087 edges, of which 60 were true interactions but 1026 false associations (i.e. environmentally-driven or due to chance). Applying each method individually, we detected a moderate to high number of environmentally-driven edges—87% Sign Pattern and Overlap, 67% Interaction Information, and 44% Data Processing Inequality. Combining these methods in an intersection approach resulted in retaining more interactions, both true and false (32% of environmentally-driven associations). After validation with the simulated datasets, we applied EnDED on a marine microbial network inferred from 10 years of monthly observations of microbial-plankton abundance. The intersection combination predicted that 8.3% of the associations were environmentally-driven, while individual methods predicted 24.8% (Data Processing Inequality), 25.7% (Interaction Information), and up to 84.6% (Sign Pattern as well as Overlap). The fraction of environmentally-driven edges among negative microbial associations in the real network increased rapidly with the number of environmental factors.

**Conclusions:** To reach accurate hypotheses about ecological interactions, it is important to determine, quantify, and remove environmentally-driven associations in marine microbial association networks. For that, EnDED offers up to four individual methods as well as their combination. However, especially for the intersection combination, we suggest using EnDED with other strategies to reduce the number of false associations and consequently the number of potential interaction hypotheses.

## Background

### Association networks to generate microbial interaction hypotheses

There is a myriad of microorganisms on Earth; current estimates indicate ≈10^12^ microbial species (Locey & Lennon, 2016), and ≈10^30^ microbial cells (Whitman *et al.*, 1998; Kallmeyer *et al.*, 2012). Microorganisms have crucial roles in the biosphere by contributing to global biogeochemical cycles (Falkowski *et al.*, 2008) and underpinning diverse food webs. The importance of microbes for the functioning of ecosystems cannot be understood without considering their ecological interactions (DeLong, 2009; Krabberød *et al.*, 2017). These allow transferring carbon and energy to upper trophic levels, and the recycling of nutrients and energy (Worden *et al.*, 2015). Furthermore, ecological interactions influence microbial community turnover and composition. These interactions include win-win (e.g. mutual cross-feeding and cooperation), win-loss (e.g. predator-prey and host-parasite), and loss-loss (e.g. resource competition) relationships (Faust & Raes, 2012). Although microbial communities are highly interconnected (Layeghifard *et al.*, 2017), our knowledge about ecological interactions in the microbial world is still limited (Krabberød *et al.*, 2017; Bjorbækmo *et al.*, 2019).

Previous studies have shown relationships between a restricted number of microorganisms. However, we need a large number of interactions to understand the functioning of complex ecosystems. This is challenging, in part, due to the vast number of possible interactions—given n microorganisms, there are 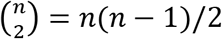 potential pairwise interactions. Thus, it is unfeasible to test them experimentally within a reasonable amount of time and cost. The problem of having a large number of potential interactions can be partially circumvented with omics technologies coupled to network analyses.

Omics can identify and quantify a large number of microorganisms from a given sample. Typically, the relative abundance for each identified organism per sample is estimated. There are multiple methods to determine associations (normally based on correlations) between microorganisms using their abundances (e.g. eLSA (Xia *et al.*, 2011, 2013), CoNet (Faust & Raes, 2016), SPIEC-EASI (Kurtz *et al.*, 2015), or FlashWeave (Tackmann *et al.*, 2019)). These abundance-based associations compose a network, where nodes represent microorganisms and edges represent either co-presence (positive association) or mutual exclusion (negative association) relationships, which constitute microbial interaction hypotheses.

### Challenges in using networks as a representation of the microbial ecosystem

Although networks play an essential role in understanding complex systems, microbial ecological networks are not yet as developed in terms of inference and biological interpretation (Lv *et al.*, 2019). Network inference from -omics data is difficult (Li *et al.*, 2016; Layeghifard *et al.*, 2017) because of both technical and interpretation challenges. One challenge is the compositional nature of the data produced by DNA sequencers (Gloor *et al.*, 2017). There are several network tools (Li *et al.*, 2016) that consider this, e.g., SPIEC-EASI (Kurtz *et al.*, 2015). Other difficulties include data based on a small number of samples relative to the number of microorganisms they contain, i.e., a low sample-to-microorganisms ratio; plus sparse data—too many zeros in the dataset that can wrongly associate microorganisms (Aitchison, 1981). A zero indicates either the absence of a microorganism (structural zero), or an insufficient detection level or sequencing depth (sampling zero). Thus, we should remove microorganisms appearing in just a few samples.

Interpretation of association networks is challenging because they are not equivalent to ecological networks. Edges in ecological networks represent observed ecological interactions between different microorganisms like parasitism or competition (Xiao *et al.*, 2017). Ecological networks are directed graphs, where the directed edges (arcs) point from a start node (source) to an end node (target). In contrast, association networks are undirected. Although association networks provide ecological insight, they do not necessarily encode causal relationships or observed ecological interactions. Unless edges are verified with experiments or additional information, one should be careful when attributing biological meaning to network properties (Röttjers & Faust, 2018). In addition, networks with too many edges (dense networks or hairballs) make interpretation more challenging. We can reduce network density when lowering the corrected *p*-value for inferred edges (Weiss *et al.*, 2016), or increasing the cut-off for other criteria such as the association strength, prevalence, or abundance filtering (Röttjers & Faust, 2018). Another strategy is agglomeration using taxonomic or ecological (functional) groupings (Lima-Mendez *et al.*, 2015).

The interpretation challenge addressed in this study are indirect dependencies (associations) caused by environmental factors. For most microbial association networks, an edge indicates one of the following three alternatives:

1. ecological interaction between two microorganisms,
2. similar or contrary dependence (i.e., preference) to environmental factor/s or a third microorganisms,
3. association by chance.

Indirect associations occur when two microorganisms are both dependent on an abiotic environmental factor (e.g., same nutrients and temperature requirements) or biotic factor (e.g., same prey or predator), but do not interact with one another. Here, indirect association describes the computational effect of indirect dependencies, and observing an association when in fact there is none.

### Removing indirect dependencies including environmental effects

To distinguish between direct and indirect interactions, several network construction tools use a probabilistic graphical model (Kurtz *et al.*, 2015; Yang *et al.*, 2017), e.g. SPIEC-EASI (Kurtz *et al.*, 2015, 2019), miic (Verny *et al.*, 2017), or FlashWeave (Tackmann *et al.*, 2019). FlashWeave can also integrate metadata to avoid indirect associations driven by environmental factors but currently does not support missing data. The tool ARACNE (Margolin *et al.*, 2006) aims to eliminate indirect associations by using an information theoretic property (the *Data Processing Inequality*, DPI, in Methods). The extension TimeDelay-ARACNE (Zoppoli *et al.*, 2010) tries to extract dependencies between different times. Another approach including time-delay is implemented in the tool MIDER (Villaverde *et al.*, 2014), which combines mutual information-based distances and entropy reduction to detect indirect interactions (*Mutual Information*, MI, in Methods). PREMER (Villaverde *et al.*, 2018), a successor of MIDER, allows to include previous knowledge, e.g., known non-existent associations.

There are also several prior network construction approaches to reduce indirect associations, e.g., a high prevalence filter that preserves microorganisms present in many samples (Pascual-García *et al.*, 2014). However, this will keep generalist while removing specialist. Another approach divides datasets displaying a great environmental heterogeneity into sub datasets of similar environmental conditions (Röttjers & Faust, 2018). For example, a previous work (Mandakovic *et al.*, 2018) constructed two networks representing bacterial soil communities from two different sections of a pH, temperature, and humidity gradient. Another work (Lima-Mendez *et al.*, 2015) constructed ocean depth-specific networks to account for environmental differences between the surface layer and the deep chlorophyll maximum layer. In addition to dividing samples, an algorithm aiming to correct for habitat filtering effects (Brisson *et al.*, 2019), subtracts, for a given habitat, the mean abundance from each microorganisms within each sample. However, this approach is limited to the identified habitat groups that should have a similar sample size.

In contrast, there are methods accounting for indirect dependencies after network construction. For instance, global silencing, (Barzel & Barabási, 2013) and network deconvolution (Feizi *et al.*, 2013) aim to recover true direct associations from observed correlations. Both techniques are sensitive to missing variables (Alipanahi & Frey, 2013). Another method, called *Sign Pattern*, SP, uses environmental triplets (Lima-Mendez *et al.*, 2015). An environmental triplet contains two microorganisms and one environmental factor, which are associated to each other. SP combines the signs of association scores (positive or negative) to determine if a microbial association should be classified as indirect (SP in Methods). Its major drawback is edge removal where microorganisms with similar environmental preference interact. Along SP and network deconvolution, the *Interaction Information*, II, was applied in (Lima-Mendez *et al.*, 2015). Within an environmental triplet, the II method aims to indicate whether an edge is due entirely to shared environmental preferences (II<0) or whether environmental preferences and true interactions are entangled (II>0). However, II cannot determine which associations in a triplet is indirect (II in Methods). Here, we study several indirect edge detection methods: SP, *Overlap*, (OL, developed here), II, DPI, and their combination.

### EnDED is an implementation of four methods and their combination

This article presents EnDED, which implements four approaches, and their combination, to indicate environmentally-driven (indirect) associations in microbial networks. The four methods are: Sign Pattern (Lima-Mendez *et al.*, 2015), Overlap (developed here), Interaction Information (Lima-Mendez *et al.*, 2015; Ghassami & Kiyavash, 2017), and Data Processing Inequality (Cover & Thomas, 2001; Margolin *et al.*, 2006). SP requires an association score that represents co-occurrence when it is positive, and mutual-exclusion when it is negative. OL requires temporal data with a known start and end of the association to determine whether the microbial association occurs in a time window when both microorganisms are associated to the same environmental factor. The II method indicates the existence of one indirect dependency between three components that are associated with each other. The DPI method states that the association with the smallest mutual information is the indirect association. Here, we evaluate each method and their combination on how well they detect environmentally-driven associations on association networks from simulated data including two environmental factors. Combining methods in an intersection approach retains more true interactions than each method on its own. A union approach was discarded because it would have retained the smallest number of true interactions. We are able to disentangle and filter environmentally-driven edges from microbial association networks (0.95-0.96 in positive predictive value and 0.35-0.83 in accuracy). We also applied EnDED to disentangle and filter environmentally-driven edges from a real marine microbial association network based on ten years of monthly sampling including ten environmental factors. EnDED contributed to both, generating more reliable hypotheses on microbial interactions, and facilitating network analysis by removing edges from dense “hairball” networks. EnDED is publicly available (Deutschmann, 2019).

## Results

### Simulated data

To evaluate EnDED’s performance in removing environmentally-driven associations, we simulated 1000 abundance time-series datasets with 50 microorganisms and known true interactions between them. We obtained another 1000 datasets with noise (hereafter dwn). We constructed the networks (hereafter simulated networks) with the tool eLSA (Xia *et al.*, 2011, 2013) (see methods). The simulated networks contained on average (computed as the median) 50 nodes and 1087 edges (1063 dwn), of which 60 (59 dwn) were true interactions (edges present in the inferred and true network) and 1026 (1005 dwn) false associations (edges present in the inferred but absent in the true network). Networks inferred from simulated data without noise contained on average one more true interaction but also 21 more false interactions than the networks inferred from simulated data with noise.

A simple approach to discriminate true interactions (desired) from false associations (undesired) would be to use a threshold for the association strength, which could be suitable if the values for true interactions and false associations are i) following different distributions, and ii) the distributions are mainly non-overlapping. We tested the former requirement with a two-sample Kolmogorov-Smirnov test with the R (R Core Team, 2019) function *ks.test*. Using a 95% (99%, 99.9%) confidence level, the distributions were significantly different for 358 (192, 66) simulated datasets and 355 (173, 68) simulated datasets with noise, which is slightly more than one third of them. This indicates that an association strength cut-off is unsuitable to separate true interactions from false associations. More sophisticated approaches than a simple threshold include the methods implemented in EnDED: SP, OL, II, DPI, and their combination.

Combining the methods in an intersection approach (hereafter referred to as intersection combination), we classified on average 348 (228 dwn), that is 32% (22% dwn) of the associations, to be environmentally-driven. The number of correctly detected false associations was on average 332 (219 dwn), i.e., 96% of the removed edges. The resulting networks contained on average 737 (828 dwn) edges. When each method was individually applied more edges were removed: 87% (86% dwn) for SP and OL, 67% (60% dwn) for II, and 44% (32% dwn) for DPI. The fraction of correctly removed edges for individual methods was on average 95%. Comparing the methods on correctly detected false associations, the greatest agreement was observed between SP and OL, whereas DPI appeared to be the most conservative in not agreeing with other methods and, subsequently, reducing the number of detected edges in the intersection combination approach (Supplementary Table S1).Individual methods removed more edges from the network than the intersection combination, where all methods must agree. However, a method’s performance is not solely determined by the number of removed edges.

To evaluate the removal of environmentally-driven edges, we scored the different approaches based on five evaluation measurements (see Methods): the true positive rate, TPR, true negative rate, TNR, false positive rate, FPR, positive predicted value, PPV, and accuracy, ACC, (Figure 1 and Supplementary Table S2). In order to determine these measurements, we first determined true and false positives, as well as true and false negatives. A true positive is a false association in the network that is correctly removed by a method, and a false negative is a false association that is incorrectly not removed. A false positive is a true interaction in the network that is incorrectly removed by a method, and a true negative is a true interaction that correctly is not removed by a method. The ideal method maximizes true positives and true negatives and minimizes false positives and false negatives.

**Figure 1:**
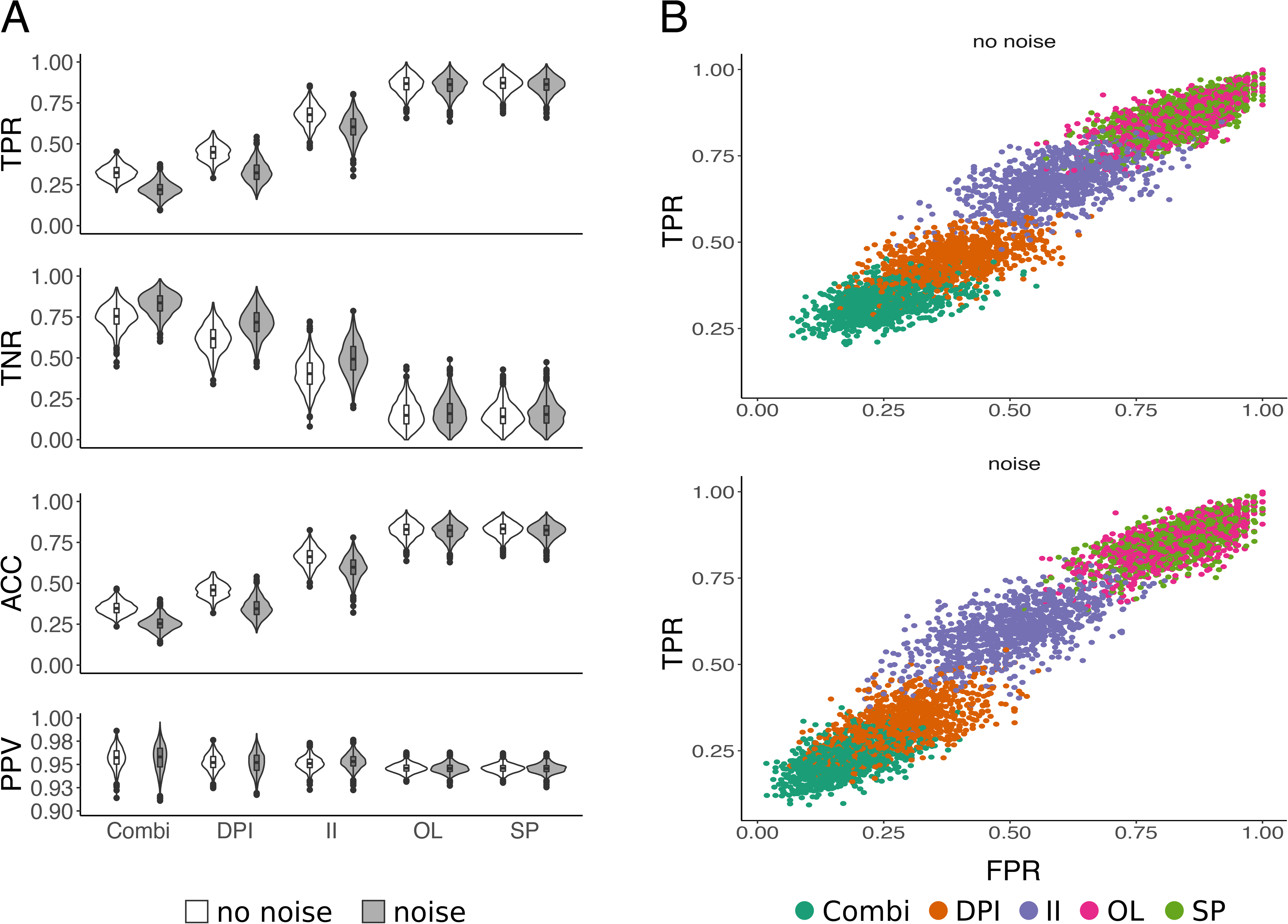
Evaluation of EnDED: intersection combination and individual methods on simulated networks. Using 1000 simulated networks, and 1000 simulated networks incorporating noise, we evaluated EnDED’s performance. Plot A) displays the evaluation measurements true positive rate (TRP), true negative rate (TNR), accuracy (ACC), and positive predictive value (PPV) for each individual method, i.e., Sign Pattern (SP), Overlap (OL), Interaction Information (II), and Data Processing Inequality (DPI), as well as the intersection combination (Combi). SP and OL perform best according to TRP and ACC, while the intersection combination performs best according to TNR. All methods performed well according to PPV. The intersection combination, DPI and II performed better on noisy data according to TNR because less edges were removed along with less true interactions. Plot B) displays the ROC curve for each environmentally-driven edge detection method as well as their intersection combination.

The intersection combination under-performed compared to each individual method, SP and OL perform best, and II performs better than DPI according to TPR, FPR and ACC (Figure 1). However, applying each method individually has the drawback of removing more true interactions. On average there are 60 (59 dwn) true interactions in the simulated networks. The individual methods removed 86% (85% dwn) (SP), 85% (84% dwn) (OL), 60% (51% dwn) (II), and 38% (28% dwn) (DPI). Therefore, although the intersection combination removed fewer edges, it outperformed the others according to the TNR because it eliminated fewer of the true interactions, 25% (16% dwn). All methods had high PPV values with half of all measured PPV above ≈0.95. According to PPV, intersection combination performed best and SP and OL performed worst (Figure 1).

### Real data

After testing EnDED’s performance on simulated networks, we applied it to a real microbial association network, which was constructed from 10 years of monthly samples from January 2004 to December 2013 at the Blanes Bay Microbial Observatory (BBMO) (Gasol *et al.*, 2016). These samples included bacteria and eukaryotes of two size-fractions: picoplankton (0.2-3 μm) and nanoplankton (3-20 μm). We estimated community composition via metabarcoding of the 16S and 18S rRNA gene, and inferred an association network, hereafter referred to as BBMO network (see Methods). The BBMO network contained 762 nodes including 754 ASVs and eight of the ten available environmental factors, and 30498 edges including 29820 microbial edges and 607 edges between a microorganism and an environmental factor. The network contained more positive (24458, 82.0%) than negative (5362, 18.0%) microbial associations (Figure 2).

**Figure 2:**
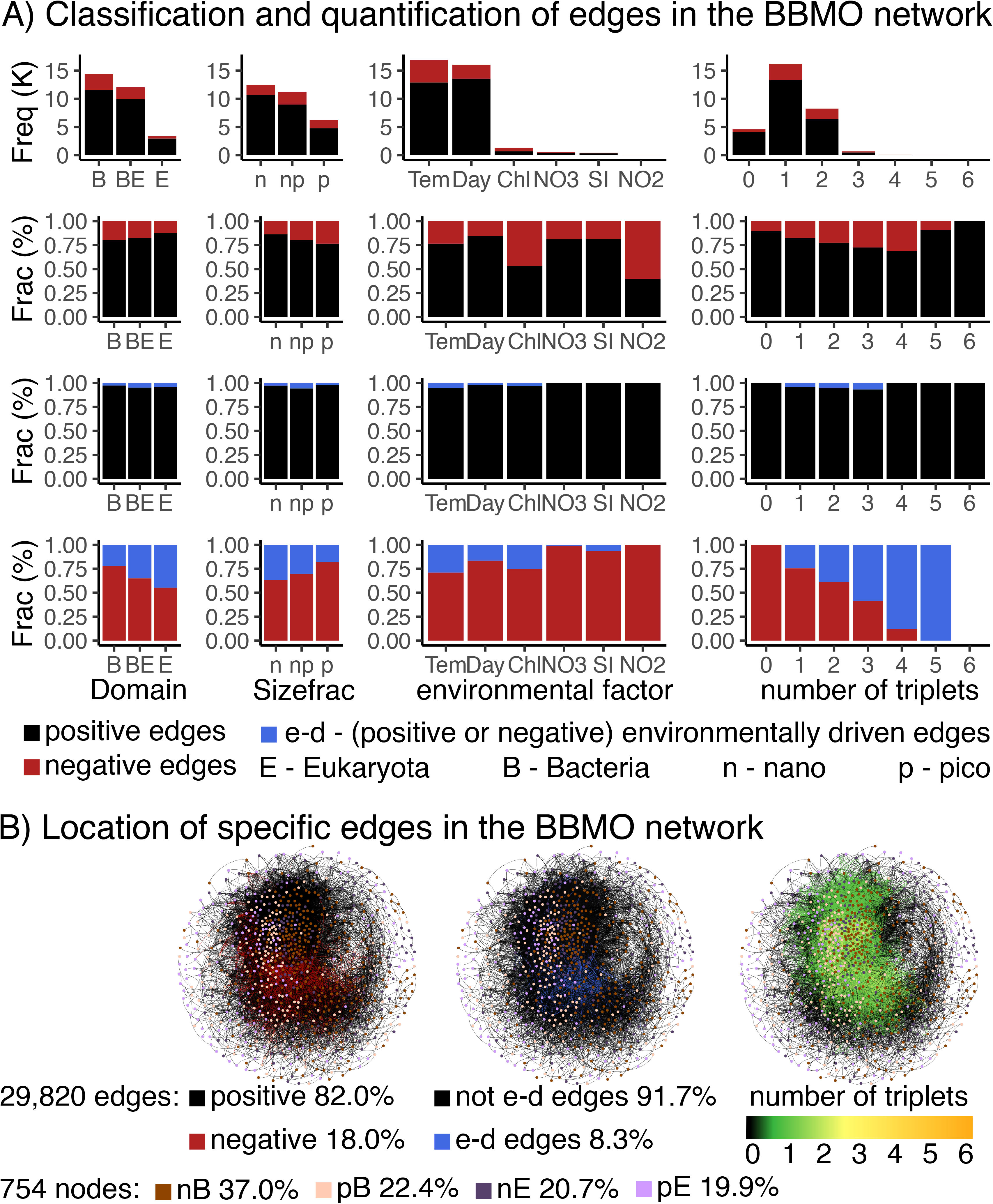
Quantification of environmentally-driven associations in the BBMO network. For A) the first column shows the number and fraction of microbial associations divided by domain: Bacteria-Bacteria associations (B), Bacteria-Eukaryote associations (BE), and Eukaryote-Eukaryote associations (E). The second column shows the number and fraction of associations divided by size-fractions: association within the nano size fraction (n), within the pico size fraction (p), and between these two size fractions (np). The third column shows all microbial edges connected to an environmental parameter: Temperature (Tem), Day length (Day), Chlorophyll (Chl), inorganic nutrients NO_3_^−^(NO3), SiO_2_ (Si), and NO_2_^−^(NO2). The last column shows the number and fraction of edges divided in how many triplets they have been found ranging from no triplets (0) to six triplets. The first two rows display the number and fraction of microbial associations of the BBMO network before applying EnDED. Positive associations are indicated with black, negative associations with red. The last two rows indicate in blue the fraction of environmentally-driven edges among the positive (third row) and negative (fourth row) microbial associations. B) The left network shows in black the positive and in red the negative associations. The right network shows the number of triplets a microbial edge is in ranging from one (green) to six (orange), and no triplet (black). The middle network shows in blue the environmentally-driven associations that were detected by the intersection combination of the four methods Sign Pattern, Overlap, Interaction Information, and Data Processing Inequality.

We found that 25230 (84.6%) of the network edges were in at least one and in maximum six environmental triplets (Figure 2 and Supplementary Table S3). Overall, we detected 35166 environmental triplets within the BBMO network. Of the ten considered environmental factors, PO_4_^3-^ and salinity were not associated to any microorganism in the network, and turbidity and NH_4_^+^ were not found within a triplet. Thus, six environmental factors remained: Temperature (1831 environmentally-driven edges were removed due to Temperature) and day length (652 removed edges) were the top two environmental factors affecting microbial associations, followed by total chlorophyll (175), SiO_2_ (5) and NO_3_^−^ (1); no edge was removed due to NO_2_^−^.

The intersection combination removed 2488 (≈8.3%) associations from the BBMO network. We classified and quantified these indirect edges according to the domain of the nodes (bacteria - eukaryotes, nanoplankton – picoplankton), environmental factor, and the number of triplets a microbial edge was in (Figure 2 and Supplementary Table S4). Compared to the intersection combination, each method individually removed more edges: 84.6% (SP and OL removing all microbial edges present in a triplet), 25.7% (II), and 24.8% (DPI); that is, removal was 3 to 10 times larger.

We also determined for each association the Jaccard index, which indicates how often two microorganisms appear together in the dataset. We assumed that two microbes that appear together < 50% of the time are less likely to have true contemporary ecological interactions and the corresponding association is more likely to be false. We found that only 27.7% of the indirect associations had a Jaccard index above 0.5 compared to 61.1% of the associations that were not indirect. This discrepancy was bigger for negative edges, with 1.2% above and 98.8% below 0.5 (Table 1). The fact that over 72.3% of environmentally-driven associations had a Jaccard index equal or below 0.5 strengthened the decision of their removal.

**Table 1:** Jaccard index of edges. The BBMO network before applying EnDED contained 29820 edges of which 2488 (8.3%) were environmentally-driven (indirect). Considering the Jaccard index for these indirect edges, 688 (27.7% of indirect edges) score above 50%, and 1800 (72.3%) score below or equal to 50%. In contrast, 61.1% of edges not considered as indirect have a Jaccard index above 50%, and 38.9% of all not indirect edges have a Jaccard index equal or below 50%.

|  | All edges | Jaccard index>50 | Jaccard index≤50 |
| --- | --- | --- | --- |
| BBMO network | 29 820 (100%) | 17 383 (58.3%) | 12 437 (41.7%) |
| positive edges | 24 458 (82.0%) | 17 212 (70.4%) | 7 246 (29.6%) |
| negative edges | 5 362 (18.0%) | 171 (3.2%) | 5 191 (96.8%) |
| indirect (intersection) | 2 488 (8.3%) | 688 (27.7%) | 1 800 (72.3%) |
| positive + indirect (intersection) | 934 (3.1%) | 670 (71.7%) | 264 (28.3%) |
| negative + indirect (intersection) | 1 554 (5.2%) | 18 (1.2%) | 1 536 (98.8%) |
| not indirect (all) | 27 332 (91.7%) | 16 695 (61.1%) | 10 637 (38.9%) |
| not indirect (min 1 triplet) | 22 742 (76.3%) | 14 242 (62.6%) | 8 500 (37.4%) |
| not indirect (no triplet) | 4 590 (15.4%) | 2 453 (53.4%) | 2 137 (46.6%) |
| Sign Pattern | 25 230 (84.6%) | 14 930 (59.2%) | 10 300 (40.8%) |
| Overlap | 25 230 (84.6%) | 14 930 (59.2%) | 10 300 (40.8%) |
| Interaction Information | 7 672 (25.7%) | 4 962 (64.7%) | 2 710 (35.3%) |
| Data Processing Inequality | 7 394 (24.8%) | 1 862 (25.2%) | 5 532 (74.8%) |

The intersection combination removed more negative than positive edges, 1554 and 934, respectively (Figure 2). However, there were 20334 positive and 4896 negative microbial associations that were found in at least one environmental triplet, so the method removed 31.7% of the negative and only 4.6% of the positive edges. If we randomly removed 2488 edges, we would expect 18.0 % to be negative (i.e. 448) and 82.0 % of them to be positive (i.e. 2040). If we restrict these calculations to the 25230 microbial associations that were found in at least one environmental triplet, with 20334 of them being positive and 4896 being negative, we would expect to remove 19.4% (i.e. 483) of negative and 80.6% (i.e. 2005) of positive edges. The probability of randomly removing less positive than negative associations is nearly zero, since it follows a multivariate hypergeometric distribution:

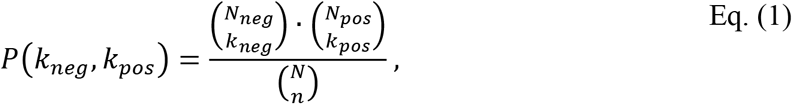

where *N*_*pos*_ and *N*_*neg*_ are the number of positive and negative associations in the network, respectively, *k*_*pos*_ is the number of removed positive and *k*_*neg*_ the removed negative associations from the network, *N* is the number of associations in the network, and *n* is the number of removed associations from the network. The removal of more negative edges through intersection combination indicates that this removal was not random or, in other words, that negative associations are more likely to represent environmentally-driven edges.

To evaluate the performance of EnDED on the BBMO network, we considered interactions described in literature and collected in the Protist Interaction Database (PIDA) (Bjorbækmo *et al.*, 2019). Studies typically compare the associations of a network to those reported in the literature at the genus level (Lima-Mendez *et al.*, 2015). The ambiguity in taxonomic classification and the large number of edges challenged this comparison. Thus, we implemented a function to compare strings and match the taxonomic classification of a microorganism in the BBMO network to those in the scientific literature (PIDA). We found that only 29 (0.1%) associations were supported by interactions described in the literature (Table 2). That is, 99.9% of associations in the BBMO network (before applying EnDED) could not be used to evaluate EnDED’s performance. These 29 associations describe eight unique interactions between eight microorganisms, and 18 edges were in an environmental triplet to which each method as well as their combination were applied (summary in Table 2). Ideally none of these described associations should be removed by EnDED. Yet, the intersection combination removed five associations (Table 2). In contrast and even worse, SP and OL removed all 18 edges, II eight and DPI nine edges. The additionally removed edges by individual methods are associations between a diatom (*Thalassiosira*) and an unknown *Flavobacteriia*. Considering only the genus level, there were 171 unique genera in the BBMO network, and 700 in PIDA, combined there were 837 microbial genera, and 34 genera in both. Thus, 19.9% of the microbial genera found in the BBMO network were also in PIDA, and 4.9% of the genera found in PIDA were also found in the BBMO network.

**Table 2:** Interactions found in the BBMO network that have been reported in the literature. The table mentions whether or not the associations were removed or kept by EnDED via the combination interaction approach. For example, the association between the ASVs classified as Dia. Thalassiosira and ASVs classified as F. unknown Flavobacteriia has been found 17 times in the network: 4 were removed and 13 were kept.

| Microorganisms | EnDED | ID in PIDA |
| --- | --- | --- |
| Included in 1, 2, 3, or 4 triplets |  |  |
| Dia. Thalassiosira - Dino. Heterocapsa | 1 removed | 1665 |
| Dia. Thalassiosira - F. unknown Flavobacteriia | 4 removed<br>13 kept | 2199 |
| Not included in a triplet |  |  |
| Dino. Heterocapsa - Dino. Prorocentrum | 1 kept | 1501, 1511 |
| Dino. Gyrodinium - Dino. Heterocapsa | 1 kept | 1313, 1314, 1780, 1783 |
| Dino. Prorocentrum - Dino. Gymnodinium | 2 kept | 1499 |
| Dino. Prorocentrum - Dino. Prorocentrum | 4 kept | 1509, 1510 |
| Dino. Prorocentrum - Dino. Scrippsiella | 2 kept | 1513 |
| F. unknown Flavobacteriia - Dia. Pseudo-nitzschia | 1 kept | 2196 |

Abbreviations indicate Dia - Diatomea; Dino - Dinoflagellata; C - Ciliophora; F - Flavobacteriia; ID in PIDA refers to the number PIDA gave to an interaction described in the literature.

## Discussion

### Using EnDED to disentangle environmental effects in microbial association networks

EnDED makes several indirect-edge removal techniques accessible to microbial ecologists without requiring previous programming experience. These techniques can be used individually or combined. In addition, this work systematically evaluated the different techniques and their combination to remove indirect edges from microbial association networks. Here, we tested only the union and intersection combination of all four methods, but other combination strategies are possible with EnDED. EnDED requires data of the environmental factors in order to predict if an association is environmentally-driven. This is a limitation, since it may be impossible to consider all environmental factors (Lv *et al.*, 2019). However, EnDED can perform well if the major environmental factors, such as, e.g., temperature and nutrient concentrations for marine microorganisms, are provided. Moreover, knowledge of microbial interactions in nature is rather limited and therefore determining the performance of EnDED for real networks is challenging and carries some degree of uncertainty. Thus, EnDED’s results should be interpreted with care.

For the simulated networks, we found that each method individually removed on average a moderate to high number of edges. The intersection combination removed fewer edges but kept more true interactions. To understand the impact of the environment, Röttjers and Faust simulated an increasing environmental influence and observed a decrease in retrieving true interactions from inferred associations (Röttjers & Faust, 2018). The observation holds for several network construction methods for cross-sectional data, including CoNet (Faust *et al.*, 2012), SparCC (Friedman & Alm, 2012), SPIEC-EASI (Kurtz *et al.*, 2015), and Spearman correlations. In agreement with these findings, we observed a slight increase in retrieving true interactions when removing environmentally-driven associations in our simulation networks.

In our BBMO dataset, the intersection combination removed a modest number of the edges—a much higher fraction of negative than positive edges. We argue that several negative associations are probably due to different environmental preference (different niches) of microorganisms. The Jaccard index representing a level of microbial co-occurrence, scored equal or below 50% for most negative associations. These may partially represent microorganisms adapted to different seasons. Previous work on the eukaryotic pico- and nano-plankton at the BBMO, using the same basal 10-year dataset used here, indicated a strong seasonality at the community level (Giner *et al.*, 2019).

### Comparisons of indirect edge detection on other datasets

In our BBMO network we found that the majority (84.6%) of the microbial edges was within at least one environmental triplet. This was 2.6 times higher than what was found for an association network inferred from data considering microorganisms and small metazoans from two ocean depths across 68 stations around the world and various size fractions (hereafter global interactome) (Lima-Mendez *et al.*, 2015). This global interactome contains 29912 (32.3%) edges that were within at least one environmental triplet (Lima-Mendez *et al.*, 2015). In the previous study, 29900 edges in the global interactome (≈100% of triplets and 32% of all edges) were attributed to environmental factors by SP, similarly to this study as SP removed all edges within triplets in the BBMO network. II indicated 11043 environmentally-driven edges in the global interactome (≈37% of triplets and 12% of all edges) with *p*-value below 0.05 in a permutation test with 500 iterations. In comparison, II removed a higher fraction of edges in the BBMO network when considering all edges (25.7%), but less when considering within the triplets (30.4%). Network deconvolution suggested 22439 environmentally-driven edges (≈75% of triplets and 24% of all edges) within the global interactome, and the three methods agreed for 8209 edges (≈27% of triplets and 8.9% of all edges). In comparison, we detected slightly less environmentally-driven associations for the BBMO network (8.3% of all edges). These differences suggest that a higher environmental heterogeneity in the dataset may induce more indirect edges. Also, the effects of indirect dependencies may depend on dataset type (e.g., temporal vs. spatial). These possible differences and their effect on environmentally-driven edges should be further investigated.

Using II for the BBMO network, we identified a moderate number of environmentally-driven associations. DPI also identified a moderate number (24.8%, 29.3% when considering only triplets), whereas SP or OL identified a ubiquitous number of environmentally-driven edges (84.6%, 100% when considering only triplets). This indicates that SP and OL are strict and should be used in combination with other methods in an intersection approach.

In another study, the tool FlashWeave (Tackmann *et al.*, 2019) predicted direct microbial interactions in the human microbiome using the Human Microbiome Project (HMP) dataset, including heterogeneous microbial abundance data of 68818 samples (The Human Microbiome Project Consortium: Huttenhower *et al.*, 2012). The inferred networks (with and without metadata) were sparser than our networks. The network with metadata contained 10.7% fewer associations compared to the network without metadata, slightly more than in our results from BBMO.

### Factors causing indirect microbial associations

From the simulated networks, we found that using the intersection combination instead of each method individually, we maintained more true interactions at the cost of more false associations in the network—more when considering simulated networks including noise. Comparing our simulated network against the BBMO network, the intersection combination classified a higher number of edges as environmentally-driven in the simulated networks 32% (22% dwn) than in the BBMO network (8.3%). For the simulated data, we previously knew the environmental factor influencing pairwise microbial associations. For the BBMO data, we used ten available environmental factors, but not all factors that could affect microbial dynamics. Even though the most important factors influencing microbial seasonal dynamics at BBMO were considered (Giner *et al.*, 2019), there are several factors that were not measured and that could generate indirect edges. The indirect edges associated to these factors were not detected in our analyses. Similarly, indirect edges associated to biotic interactions (e.g., two bacteria sharing a positive edge as they are symbionts in the same protists) were not considered. Future sampling for microbial interaction research should expand metadata collection in order to detect (more) abiotic and biotic factors that could generate indirect edges.

While temperature and day length (hours of light) were the top two environmental factors affecting microbial associations in the BBMO network, the most important environmental factors in the global interactome (Lima-Mendez *et al.*, 2015) were phosphate concentration and temperature, followed by nitrite concentration and mixed-layer depth. Although we considered PO_4_^3-^ and salinity, they were not associated to any microorganism in the network, which may reflect the low variation of these environmental factors in the studied marine site (BBMO). For instance, the standard deviation in the BBMO dataset was < 1 for PO_4_^3-^ and salinity, in contrast to the global interactome dataset (Lima-Mendez *et al.*, 2015), where it was about 20-30 when considering all samples. During the Malaspina-2010 Circumnavigation Expedition, the concentrations of trace metals were determined for 110 surface water samples (Pinedo-González *et al.*, 2015). The previous study indicates relationships between primary productivity and trace nutrients, more specifically for the Indian Ocean Cd, the Atlantic Ocean Co, Fe, Cd, Cu, V and Mo, and the Pacific Ocean Fe, Cd, and V. Thus, trace metals are further environmental factors that may play an important role in regulating oceanic primary productivity.

### Limitations of EnDED

EnDED detects and removes environmentally-driven indirect edges. However, its triplet analysis could be extended to remove indirect edges driven by taxa, as done with gene triplets (Margolin *et al.*, 2006). A recent update of the network construction tool eLSA (Xia *et al.*, 2011, 2013) permits to examine how a factor, such as a microorganism or environmental variable, mediates the association of two other factors (Ai *et al.*, 2019), which allows the study of interactions between three factors. Furthermore, triplets limit the study to first-order indirect dependencies, neglecting higher-order indirect dependencies. Such limitation was solved for the DPI method by examining associations in quadruplets, quintuplets, and sextuplets (Jang *et al.*, 2013). Implementing higher-order DPI and adjusting the other three methods to account for higher-order indirect dependencies may be promising but one needs to be aware that incorporating higher-order dependencies will also increase the risk of over-fitting. Further, all relevant (measured) environmental factors could be incorporated into the calculation of II, which would combine environmental triplets. However, we reason that such adjustments would require a larger sample size. Both II and DPI calculate MI that measures the dependence between two random variables. EnDED is limited by including one function to estimate the MI. A comparison of four different MI estimates revealed that obtaining the true value of MI is not straightforward, and minor variations of assumptions yield different estimates (Fernandes & Gloor, 2010). Lastly, the conditional mutual information, CMI, which quantifies nonlinear direct relationships among variables, can be underestimated if variables have tight associations in a network (Zhao *et al.*, 2016). The so-called part mutual information, PMI, measurement can help overcome CMI’s underestimations. Although using PMI instead of CMI looks promising, calculating PMI is computationally more demanding (Zhao *et al.*, 2016).

### Future Perspectives

In this study, we have shown that EnDED with an intersection combination approach provides less dense networks, but still with many potential interactions. We observed a trade-off comparing single methods with the combination approach (intersection combination). Although the latter kept more true interactions, it kept also more false associations. Inferring emergent properties is a key task in microbial ecology to characterize microbial ecosystems from a network-perspective. Thus, if the study aim is to explore patterns of network topology rather than single edges, inferring a network comparable to the real interaction network may be more useful than accuracy of single edges. However, investigations aiming to provide potential interaction partners may use EnDED with the intersection combination approach (e.g., (Latorre *et al.*, 2021)). Specific associations may be validated with experiments or microscopy (Lima-Mendez *et al.*, 2015; Krabberød *et al.*, 2017). However, we suggest to first further reduce the set of potential interaction hypotheses. To improve the selection of interaction hypotheses, we propose to score associations based on re-occurrence: in time, as done with microbial abundance seasonality (Giner *et al.*, 2019), or space, where an association appears in different networks based on different datasets, or different regions of the world. In a previous study using 313 samples, including seven size-fractions, four domains (Bacteria, Archaea, Eukarya, and viruses), and two depths from 68 stations across eight oceanic provinces, 14% of the 81590 predicted biotic interactions were identified as local (Lima-Mendez *et al.*, 2015). Thus, re-occurrent associations may suggest a higher likelihood that the association represents a true ecological interaction, reducing the number of interaction hypotheses to the strongest ones. Another strategy to shortlist interaction hypotheses is to incorporate additional data into the network and use a multi-layer network approach. Such data could be environmental preferences such as temperature or salinity optima, size of cells, presence of chloroplasts, or data obtained from High-Throughput Cultivation (Faust, 2019), microbial community transcriptomes that reveal metabolic pathways (McCarren *et al.*, 2010), or interactions inferred from Single-Cell genome data (Yoon *et al.*, 2011; Krabberød *et al.*, 2017).

## Conclusion

In this paper, we presented EnDED, an analysis tool to reduce the number of environmentally induced indirect edges in inferred microbial networks. Applying EnDED on simulated networks indicated that false associations, driven by environmental variables instead of true interactions, were ubiquitous. However, EnDED’s intersection combination classified a minority of associations as environmentally-driven in a real (BBMO) network. Depending on the single method used, we classified a moderate to high number of associations as environmentally-driven in the same network. Nevertheless, associations driven by environmental factors must be determined and quantified to generate more accurate insights regarding true microbial interactions. EnDED provides a step forward in this direction.

## Methods

### Simulated dataset: time series based on an adjusted generalized Lotka-Volterra model

To evaluate the performance of EnDED, we simulated a time series using an adjusted version of the standard *generalized Lotka-Volterra model*, gLV (Berry & Widder, 2014; Bashan *et al.*, 2016). The gLV can describe the dynamics of microbial communities, by including a first order approach of the microbial interactions. The model’s simplicity arises from the assumption of linear interactions, which facilitates implementation and allows fast numerical simulations. The gLV has, however, several limitations (Gonze *et al.*, 2018). For example, gLV neglects higher-order interactions and the additivity of interaction strengths is a weakness because they may be combined in different ways. Also, interactions are often assumed to be constant parameters, but a reducing level of a nutrient may weaken cross-feeding relationships. Moreover, gLV omits the influence of environmental factors, which, for example, can induce oscillations in natural communities (Benincà *et al.*, 2011). Using a model that accounts for nutrients (Kettle *et al.*, 2018) is more realistic but also more complex. More elaborate mechanistic models of microbial dynamics than gLV solve explicitly the global cycling of nutrients and are coupled to the oceanic circulation (see (Vallina *et al.*, 2019) for a review), but the added complexity can hamper understanding about the ecological interactions among microorganisms when compared to a simpler gLV approach. Thus, we chose to use a simpler extension of the gLV to account for the influence of environmental factors (Stein *et al.*, 2013; Dam *et al.*, 2016). In order to allow the growth rates to vary when the environmental variables change, environmental variables can be incorporated directly into the gLV (Dam *et al.*, 2016; Röttjers & Faust, 2018). We simulated a time series using the Klemm-Eguíluz algorithm (Klemm & Eguíluz, 2002), and an adjusted gLV. We adjusted the model by defining microbial growth rates as a function dependent on one seasonal abiotic environmental factor, and added an abiotic environmental factor in the interaction matrix. We then used the time series generated by the gLV to obtain temporal microbial abundance data. With this simulated data, we inferred a network that contained environmentally-driven associations, needed to evaluate the performance of EnDED. We repeated this procedure 1000 times to obtain a large set of simulated networks, and then used the determined abundance tables and Poisson distribution to obtain another 1000 simulated networks including noise. The addition of noise was done by randomly drawing an abundance from the Poisson distribution with λ equaling the original abundance of a specific microorganisms to a specific time.

#### Adjusting the gLV

To evaluate EnDED, we simulated a time series of microbial abundances with a gLV including true pairwise interactions between 50 microorganisms and adjusted it by incorporating two environmental factors:

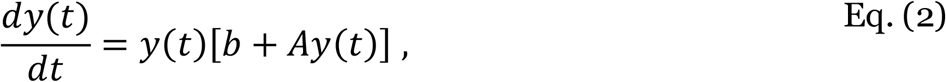

where *t* is time, *dy*(*t*)/*dt* is the rate of change of microbial abundances as a column vector, *y*(*t*) is the vector of microbial abundance at time *t*, b is the growth rate vector determined through microorganism’s specific growth rate functions that depend on an environmental factor (see equation (4)), and *A* is the interaction matrix.

#### Interaction matrix

In the interaction matrix *A*, each coefficient *a*_*ji*_ provides the linear effect that a change in the abundance of microorganism *i* has on the growth of microorganism *j* (Novak *et al.*, 2016). We simulated the interaction coefficients *a*_*ji*_ with the Klemm-Eguíluz algorithm (Klemm & Eguíluz, 2002), which generates a modular and scale-free matrix. We also set the interaction probability to 0.01, the percentage of positive coefficients to 30%, and diagonal coefficients to zero. Negative diagonal coefficients *a*_*ii*_ (i.e., the interaction of a microorganism with itself) can represent intra-specific competition and provides the carrying capacity for each microorganism, preventing its explosive growth (Haydon, 1994). We set the diagonal coefficients *a*_*ii*_ = −0.5 to avoid excessive microbial abundances in the simulations.

#### Two abiotic environmental factors

We adjusted the gLV by including two environmental factors. For simplicity, we assume no feedback between the microorganisms and the environmental factors. That is, the environmental factors affect the growth of the microorganisms but not vice-versa. The first environmental factor affects the specific growth rate of each microorganism by interacting with two of their traits: optimal environmental value for growth and tolerance range of environmental values. We simulated the environmental factor using a periodic sinusoidal function (see equation (3)), rounded to 3 digits:

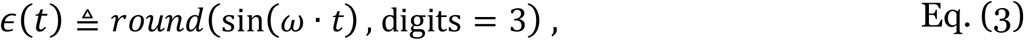

where *t* is the time axis (months), ω = (−2*π*/*T*) is the signal frequency (radians) and *T* = 12 is the signal periodicity (months); resulting in a signal phase shift of *T*/4 (months). While the first environmental factor is considered to be “external” to the microbial community, the second environmental factor is considered to be “internal”, and therefore it is included in the interaction matrix. The interaction coefficients between the microorganisms and the second environmental factor were generated by splitting the microorganisms into two groups: the second abiotic environmental factor influenced positively one half and negatively the other half of the microorganisms. We obtained the interaction coefficients from two uniform distributions defined to range between [−0.8, −0.2] and [0.2, 0.8] respectively. As the microorganisms did not influence the abiotic factor, the corresponding interaction coefficients were set to zero.

#### Species growth rate

The external seasonal abiotic environmental variable affects the growth rate, *g*, of each microorganism. This dependency is given by:

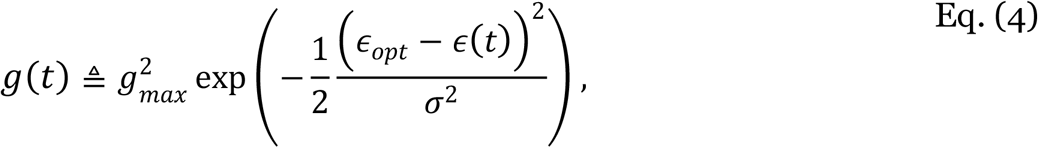

where *E*(*t*) is the environmental parameter that affects the microorganisms growth rate *g*(*t*) at time *t*, *g*_*max*_ is the microorganism’ specific maximum growth rate that determines the amplitude of the growth-rate curve, *ϵ*_*opt*_ is the microorganism’ specific optimal environmental value that determines the peak of the growth-rate curve, and *σ* is the microorganism’ specific ecological tolerance (niche width) determining the environmental range in which the microorganism grows, which determines the length (niche spread) of the growth-rate curve. We obtained the two constant parameters *g*_*max*_, and *σ* for each microorganism from a uniform distribution ranging between 0.3 and 1 to assure positive values. The values *ϵ*_*opt*_ were drawn from a uniform distribution ranging between the minimal and maximal value of the seasonal environmental factor. We defined the internal abiotic environmental factor, which is included in the interaction matrix, through the same function with *g*_*max*_ = 0.8, *ϵ*_*opt*_ = 0.5, and *σ* = 0.5. Since the growth rates depend on the environmental factor, they vary seasonally. Different microorganisms will grow better or worse at different times of the year following their environmental niches. This will lead to an asynchrony of their growth rate responses to the environment that will translate into an asynchrony of their abundances in time.

#### Initial abundances

To obtain the microbial abundances in time with the adjusted gLV, we simulated the initial microbial abundances with a stick-breaking process such that abundances add up to 1, using the function bstick (Jackson, 1993; Legendre & Legendre, 2012), and the package vegan (Oksanen *et al.*, 2019). We generated uneven initial microbial abundances without introducing zeros and set the initial value for the internal abiotic environmental factor included in the interaction matrix to 0.001.

#### Species abundances in time

Once we have set the initial conditions, we simulated microbial abundances over time by solving the equations given in the adjusted gLV (see equation (2)). Start time was 0, end time 49.5, and sample resolution 0.5 resulting in 100 samples. We used the solver function lsoda (Soetaert *et al.*, 2010). The simulated abundances in time were used to construct an association network, which is referred to as the simulated network.

### Real dataset: Blanes Bay Microbial Observatory (BBMO) time series

#### Microbial abundances

Surface water (*≈* 1m depth) was sampled monthly from January 2004 to December 2013, at the BBMO in the North-Western Mediterranean Sea (41°40′N 2°48′E) (Gasol *et al.*, 2016). About 6L of seawater were filtered and separated into picoplankton (0.2-3 μm) and nanoplankton (3-20 μm), as described in (Giner *et al.*, 2019). The DNA was extracted using a phenol-chloroform standard method (Schauer *et al.*, 2003), which has been modified by using Amicon units (Millipore) for purification.

Next, community DNA was extracted, and the 18S ribosomal RNA-gene (V4 region) was amplified in (Giner *et al.*, 2019) using the primer pair TAReukFWD1 and TAReukREV3 (Stoeck *et al.*, 2010). The 16S ribosomal RNA-gene (V4 region) was also amplified from the same DNA extracts using the primers Bakt 341F (Herlemann *et al.*, 2011) and 806R (Apprill *et al.*, 2015). Amplicons were sequenced in a MiSeq platform (2×250bp) at the sequencing service RTL Genomics in Lubbock, Texas. Read quality control, trimming, and inference of Operational Taxonomic Units (OTUs) as Amplicon Sequence Variants (ASV) was made with DADA2 v1.10.1 (Callahan *et al.*, 2016) with the maximum number of expected errors (MaxEE), set to 2 and 4 for the forward and reverse reads, respectively.

ASV sequence abundance tables were obtained for both microbial eukaryotes and prokaryotes. We subsampled both tables to the lowest sequencing depth of 4907 reads, with the rrarefy function from the Vegan package in R (Oksanen *et al.*, 2019), v2.4-2. We excluded 29 nanoplankton samples (March 2004, February 2005, and May 2010 to July 2012) featuring suboptimal amplicon sequencing. In these, we estimated microbial abundances using seasonally aware missing value imputation by weighted moving average for time series as implemented in the R package imputeTS (Moritz & Gatscha, 2017), v2.8.

Dislodging cells or particles and filter clogging can bias the collection of DNA in either small or large organismal size fractions. To reduce the bias, we divided the sequence abundance sum of the nanoplankton by the picoplankton for each ASV appearing in both size fractions and set the picoplankton abundances to zero if the ratio exceeded 2. Likewise, we set the nanoplankton abundances to zero if the ratio was below 0.5.

#### Taxonomic classification

The taxonomic classification of each ASV was inferred with the naïve Bayesian classifier method (Wang *et al.*, 2007) together with the SILVA version 132 (Quast *et al.*, 2012) database as implemented in DADA2 (Callahan *et al.*, 2016). In addition, eukaryotic microorganisms were BLASTed (Altschul *et al.*, 1990) against the Protist Ribosomal Reference database [PR2, version 4.10.0; (Guillou *et al.*, 2012)]. If the taxonomic assignment for eukaryotes disagreed between SILVA and PR2, we used the PR2 classification. We removed microorganisms identified as either Metazoa, or Streptophyta, plastids and mitochondria. In addition, we removed Archaeas since the 341F primer is not optimal for recovering this domain (McNichol *et al.*, 2021). The resulting microbial sequence abundance table contained microbial eukaryotic and bacterial ASVs. Rare ASVs were removed, i.e., we kept only ASVs present in more than 15% of the samples and with a sequence abundance sum above 100.

#### Environmental factors

We measured environmental factors that may affect the ecosystem’s dynamics. We considered a total of ten contextual abiotic and biotic variables: day length (hours of light), temperature (C°), turbidity (Secchi depth m), salinity, total cholorophyll (μg/l)^4^, and inorganic nutrients— PO_4_^3-^ (μM), NH_4_^+^ (μM), NO_2_^−^ (μM), NO_3_^−^ (μM), and SiO_2_ (μM) (Giner *et al.*, 2019). Water temperature and salinity were sampled in situ with a CTD (Conductivity, Temperature, and Depth) measuring device. Inorganic nutrients were measured with an Alliance Evolution II autoanalyzer (Grasshoff *et al.*, 2009). See (Gasol *et al.*, 2016) for specific details on how other variables were measured.

### Network construction

We constructed association networks from the simulated and the real microbial abundance tables and environmental parameters using eLSA (Xia *et al.*, 2011, 2013). We included default normalization and a z-score transformation using median and median absolute deviation. We estimated the *p*-value with a mixed approach that performs a random permutation test if the theoretical *p*-values for the comparison are below 0.05; the number of iterations was 2000. Although we are aware of time-delayed interactions and that eLSA (Xia *et al.*, 2011, 2013) could account for them, we considered our sampling interval as too large (1 month) for inferring time-delayed associations with a solid ecological basis. Thus, in our study, we focused on contemporary interactions between co-occurring microbes. For the BBMO dataset, the Bonferroni false discovery rate, *q*, was calculated for all edges from the *p*-values using the R function *p.adjust* (R Core Team, 2019). Lastly, we used a significance threshold for the *p* and *q* value of 0.001 as suggested in other works (Weiss *et al.*, 2016).

### Intersection combination of EnDED—Environmentally-Driven Edge Detection methods

EnDED includes four methods: SP, OL, II, DPI (described below) and their intersection combination (an ensemble approach of the four methods). We applied these methods to find environmentally-driven associations of microorganisms that were within an environmental triplet, as in (Lima-Mendez *et al.*, 2015). An environmental triplet is a special case of a closed triplet where one of the nodes corresponds to an environmental factor and the other two nodes correspond to microorganisms. We define the closed triplet, where there is an edge between each pair of three nodes, as *T* = {*v, w, f*} where *v* and *w* are two microorganisms, and *f* is an environmental component (see Figure 3).

**Figure 3:**
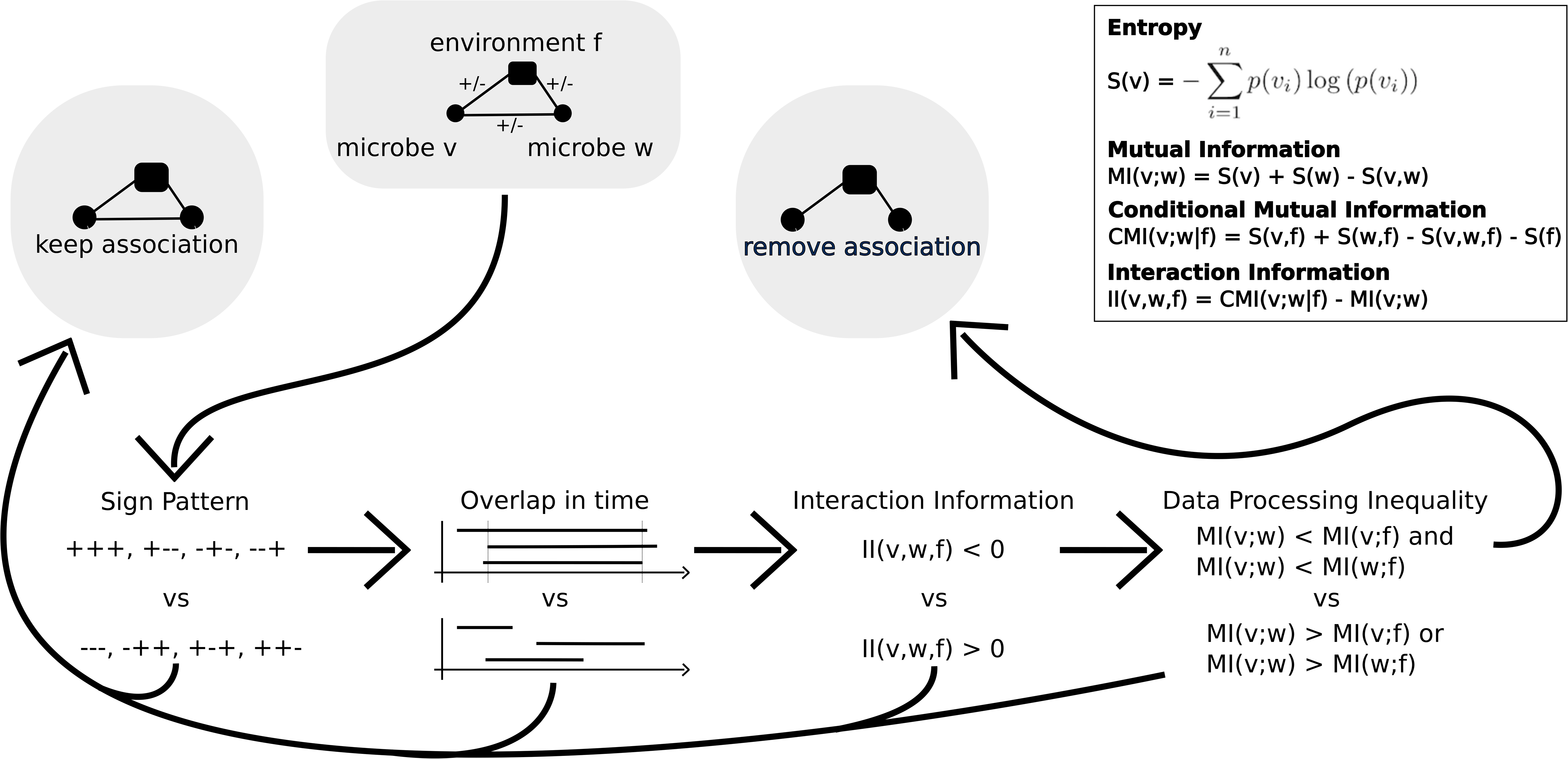
EnDED Methods Overview. EnDED is an implementation of four methods aiming to determine whether an edge between two microorganisms is indirect through the action of an environmental factor. The four methods are: Sign Pattern, Overlap, Interaction Information, and Data Processing Inequality (see Methods). Each method can be used individually or in combination. Here, we show the intersection combination approach, i.e., only if all methods classify an edge as indirect, it is removed from the network. Otherwise, the edge is classified as not indirect and kept in the network.

For the intersection combination, all four individual methods must converge to the same solution, i.e., if all methods classify the microbial edge as environmentally-driven, the edge is removed from the network. If a microbial association is within several environmental triplets, at least one of them must indicate the association as environmentally-driven. In sum, the intersection combination retains an association in the network if no triplet classifies the association as environmentally-driven.

#### Sign Pattern

The SP method (Lima-Mendez *et al.*, 2015) filters environmentally-driven edges from a network in which a positive association score indicates co-occurrence, and a negative association score indicates mutual exclusion. Let *s*_*vw*_ be the sign of the association score of the association between *v* and *w* (i.e., *s*_*vw*_ = + or *s*_*vw*_ = −). A closed triplet *T* has eight SP combinations that group into two sets (see Figure 3). If the product of the three association scores is positive, then the SP suggests that the edge between the two microorganisms is environmentally-driven. Otherwise, if the product of the three association scores is negative, SP does not suggest that the association is environmentally-driven.

#### Overlap

We have developed the OL method to support the SP for temporal data: a microbial edge should be disregarded as environmentally-driven when the associations are misaligned in time. Thus, OL requires the time when the association begins as well as how long the associations lasts, i.e., duration or length of association in time, both determined by the network construction tool eLSA (Xia *et al.*, 2011, 2013). Given an association between *v* and *w*, let 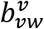 be the beginning of the association for*v*, 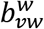 the beginning of the association for *w*, and 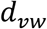 be the duration of the association between *v* and *w*. Although not used in the BBMO network, OL can consider time-delays by assuming that the beginning of the association is the minimum of the two beginnings, 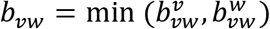, and the end of the association is the maximum, 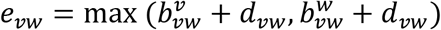. We indicate twmoicroorganisms with *v* and *w*, and the factor by *f*. The OL method calculates the overlap O of the microbial association with the two microorganism-environment associations through equation (5). As depicted in Figure 3, if *O*>60%, the microbial association is considered environmentally-driven.

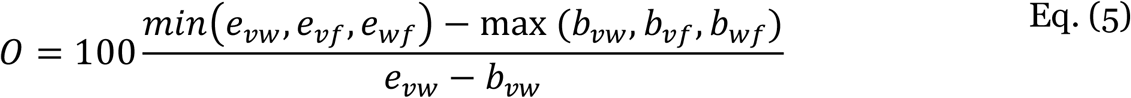

#### Mutual Information and Conditional Mutual Information

The method II employs two measurements: MI and CMI. The former is also used by DPI. Thus, before describing the methods, we first describe the two measurements. MI is a measure of the degree of statistical dependency between two variables (Margolin *et al.*, 2006). We first consider *v* = *v*_1_, … , *v*_*n*_, ***w*** = *w*_1_, … , *w*_*n*_, and ***f*** = *f*_1_, … , *f*_*n*_ as discrete random variables. The marginal probability of each discrete state (value) of the variable is denoted by *p*(*v*_*i*_) = *P* (***v*** = *v*_*i*_), the joint probability by *p*(*v*_*i*_, *w*_*j*_), and *p*(*v*_*i*_, *w*_*j*_, *f*_*k*_), and the conditional probability by *p*(*v*_*i*_|*f*_*k*_), and *p*(*v*_*i*_, *w*_*j*_|*f*_*k*_). To obtain MI, we calculate the entropy of ***v*** as

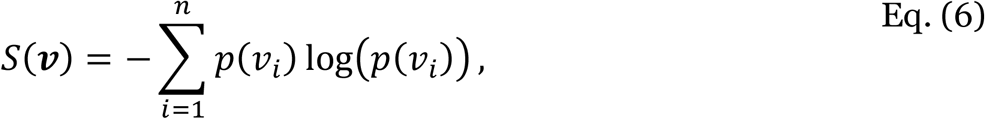

and the joint entropy of ***v*** and ***w*** as

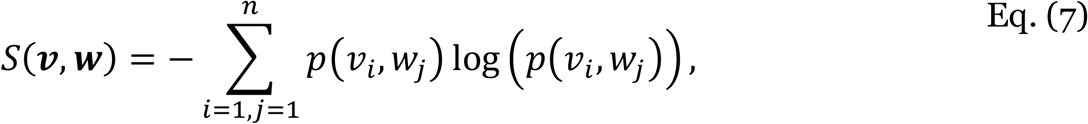

using the natural logarithm. The MI of ***v*** and ***w*** is defined through the sum of their entropies subtracted by their joint entropy:

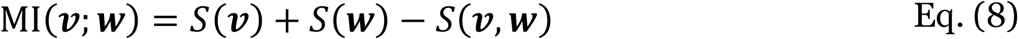

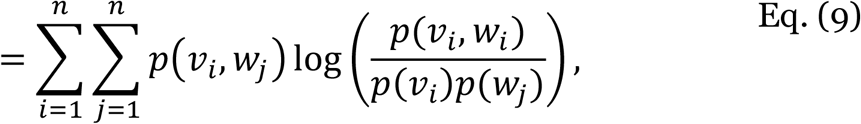

with marginal probabilities 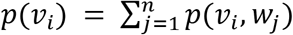, and 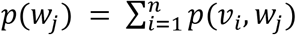.

The measurement CMI is the expected value of the MI of two random variables given a third random variable. It is defined as

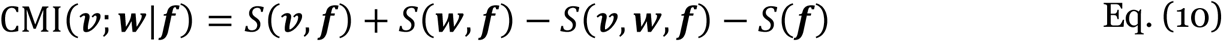

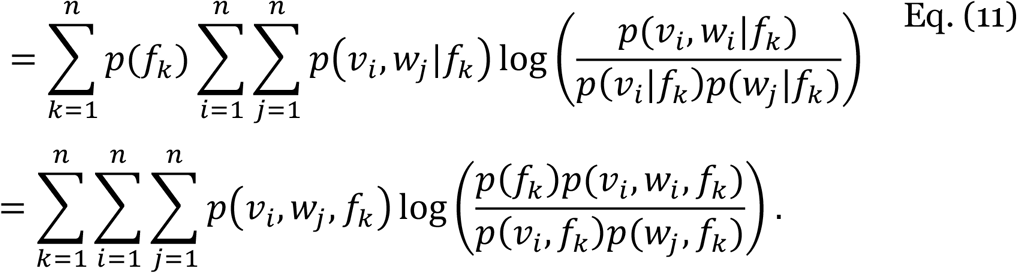

#### Interaction Information

The II is calculated with microbial abundance and environmental data. In this study, as in (Lima-Mendez *et al.*, 2015), II is computed as the difference of the CMI and MI:

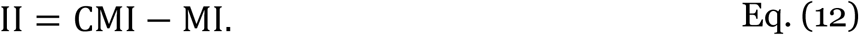

In other works (Ghassami & Kiyavash, 2017), the II is defined with a different sign convention: II = MI − CMI. In our study, if II is positive, the method suggests that the microbial association is not environmentally-driven. If II is negative, there is an environmentally-driven association within the closed triplet. However, this method cannot detect which of the three associations is indirect. In other works (Lima-Mendez *et al.*, 2015), the microbial association is assumed to be environmentally-driven if II is negative, but here we suggest to combine it with DPI (see below).

#### Significance of Interaction Information

We determined the significance of II following a strategy from (North *et al.*, 2002; Veech, 2012). We used a parameter-free permutation test and computed the *p*-value by randomizing the environmental vector ***f***. Since the MI is independent of the environmental factor and therefore remains constant, the significance of the II is the same as the CMI. Thus, we determined the significance of CMI with 1000 permutations: we randomized the environmental vector ***f*** and recalculated the CMI 1000 times, obtaining a CMI_*i*_ with *i* ∈ {1, … , 1000}. Afterwards, we quantified with *c* how many random CMI_*i*_ were at least as small as the original CMI_*i*_: *c* = |*i*: CMI_*i*_ ≤ CMI_*original*_, *i* ∈ {1, … ,1000}|. We calculated the *p*-value as

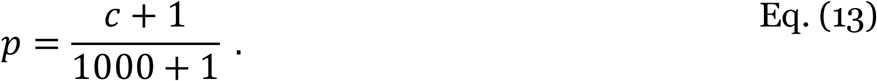

#### Data Processing Inequality

As mentioned above, the II method can detect if an indirect association exists within a triplet but cannot determine which of the three associations is indirect. Thus, we added DPI to EnDED. DPI states that if two components *v* and *w* interact only through a third component *f* (i.e., in a network *v* and *w* are connected through a path containing *f* and there is no alternative path between *v* and *w*), then the MI of *v* and *w*, MI(***v***; ***w***) is smaller than MI(***v***; ***f***) and MI(***w***; ***f***) (Cover & Thomas, 2001):

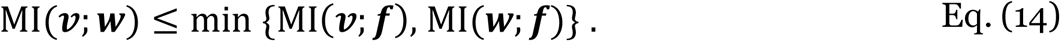

While DPI has been used in previous works on gene triplets (Margolin *et al.*, 2006), we used the DPI method for environmental triplets. We compared the MI between the two microorganisms with the MI between a microorganism and the environmental factor. If the MI between the microorganisms is the smallest, then the method suggests that the edge is environmentally-driven. This method complements the II method.

#### Equal Width Discretization

To compute the MI, CMI, and subsequently II, we discretized the abundance data and environmental parameters. EnDED uses the equal width discretization algorithm, which creates equal sized ranges (also called bins or buckets) for an abundance vector ***v*** = (*v*_1_, … , *v*_*n*_) between the lowest value (*v*_*min*_) and highest value (*v*_*max*_). It is a procedure implemented in other works (Meyer *et al.*, 2008). Given vector ***v*** of length *n* (that is the sample size) and number of bins 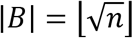, the discretized value *v*_*d*_ of variable *v* in vector ***v*** is:

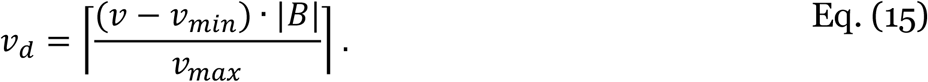

This equation assumes positive values. However, if ***v*** contains negative values, *v*_*min*_ < 0, we adjust equation (15) by substituting 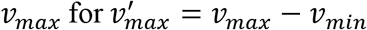. This method does not fill in missing values, and it is limited by the presence of outliers as most values would go within the same bin. We can solve this problem with a different discretization method (where bins have the same number of elements) but we have not implemented it in the current version of EnDED.

#### Applying EnDED to networks constructed from simulated and real data

We applied EnDED to association networks constructed from time series of simulated abundances and estimated microbial abundances from sequence data. The simulated networks were based on a gLV, while the real network was based on data from the BBMO. For the methods II and DPI we also included the corresponding abundance tables, and environmental factors. EnDED was run with the OL threshold of 60%. We set the significance threshold for the II score to 0.05 and used 1000 iterations.

### Evaluation of EnDED’s performance

#### Simulated network

We evaluated EnDED with the simulated interaction matrices, which revealed the number of true positives (TP), true negatives (TN), false negatives (FN), and false positives (FP) before and after removing associations that were classified as environmentally-driven. We assumed that associations not present in the interaction matrices, are environmentally-driven. We consider P as the number of all false associations, both true positive and false negative detected environmentally-driven edges: P = TP + FN, and N as the number of all true interactions, i.e., all true negative and false positive detected environmentally-driven edges: N = TN + FP. Then, we calculated the true positive rate (sensitivity), by dividing the number of true positives by the number of all real positives: TPR = (TP)/(P). Equivalently, we can also calculate the true negative rate (specificity) by dividing the number of true negatives by the number of all real negatives, TNR = (TN)/(N). The false positive rate (fall out) is the complementary to TNR, i.e. FPR = 1 − TNR. The positive predictive value (precision) can be calculated by dividing the number of true positives by the sum of all predicted positives, PPV = (TP)/(TP + FP). The accuracy is calculated by dividing the sum of true positives and true negatives by the sum of all real positives and real negatives, ACC = (TP + TN)/(P + N).

#### Real Dataset

##### Literature based database

The real network evaluation is limited since the true interactions and the microorganisms that do not interact with each other are poorly known. We assessed true interactions known in the literature based on the genus, which are compiled within the Protist Interaction Database, PIDA (Bjorbækmo *et al.*, 2019). On October 15th 2019, PIDA contained 2448 interactions. Although our dataset contains protists as well as bacteria, we were unable to evaluate interactions between bacteria through PIDA.

##### Jaccard index

In ecology, the Jaccard index (Jaccard similarity coefficient) is normally used for communities. Here, for each pair of microorganisms in the BBMO network, we computed the Jaccard index as the number of samples in which both microorganisms occur, divided by the number of samples in which at least one of the two microorganisms are present.

### Ethics approval and consent to participate

Not applicable.

### Consent for publication

Not applicable.

## Availability of data and material

EnDED is publicly available: https://github.com/InaMariaDeutschmann/EnDED. This repository contains the file “FromDataSimulationToEvaluatingEnDED.RMD”, which contains R code to generate simulated abundance tables, commands to run eLSA network construction and EnDED, as well as the command to run a C++ program (included as well) and R code used for evaluation. The repository folder BBMO data contains the BBMO abundance table, the taxonomic classification table, and the BBMO network including results of EnDED.

## Author’s contributions

IMD, GLM, JR, KF and RL designed and conceived the project. IMD performed data analysis, data simulation, and implementation of EnDED. IMD received substantial feedback on established indirect detection methods from GLM and KF, on data simulation from SMV and KF, on network construction from AKK, and on evaluation of EnDED from GLM and KF (measurements for simulation dataset) and AKK (literature based database for real dataset). RL processed the amplicon data from BBMO generating ASV tables. AKK ran the eLSA network construction tool for the BBMO dataset and IMD ran the tool for the simulation datasets. RL provided funding for the project. The original draft was written by IMD. IMD, GLM, AKK, SMV, KF and RL contributed substantially to manuscript revisions. All authors approved the final version of the manuscript.

## Acknowledgements

We thank all members of the Blanes Bay Microbial Observatory sampling team and the multiple projects funding this collaborative effort over the years. We also thank collaborators at https://www.thepapermill.eu for help with critical reading in the early stages of the manuscript. Part of the analyses have been performed at the Marbits bioinformatics core at ICM-CSIC (https://marbits.icm.csic.es).

## Supplementary Material

**Supplementary Table S1:** Comparison between methods on correctly detecting false associations. We computed the fraction (in percentage) of correctly detected false associations for each of the 1000 simulated datasets. There are only few edges that are detected by only one approach (first four rows). The most prominent groupings are highlighted in gray, e.g., SP, OL, and II agree on average on a third of edges. Combi refers to intersection combination of all four methods, SP to Sign Pattern, OL to Overlap, II to Interaction Information, and DPI to Data Processing Inequality. Less prominent groupings are aggregated with others.

| Statistic | Minimum | 1<sup>st</sup> Quartile | Median | Mean | 2<sup>nd</sup> Quartile | Maximum |
| --- | --- | --- | --- | --- | --- | --- |
| SP | 0 | 0 | 0.2 | 0.3 | 0.5 | 3.7 |
| OL | 0 | 0 | 0.1 | 0.2 | 0.3 | 2.0 |
| II | 0 | 0.7 | 1.3 | 1.4 | 2.0 | 6.0 |
| DPI | 0 | 0.1 | 0.3 | 0.4 | 0.6 | 2.6 |
| SP and OL | 4.9 | 12.2 | 14.9 | 15.0 | 17.5 | 30.0 |
| SP, OL, and II | 19.1 | 29.5 | 32.6 | 32.8 | 36.2 | 49.6 |
| SP, OL, and DPI | 2.6 | 7.1 | 8.9 | 9.1 | 10.8 | 22.1 |
| SP, OL, II, DPI, and Combi | 22.4 | 32.1 | 35.6 | 35.5 | 38.6 | 48.6 |
| other | 0.4 | 3.3 | 4.9 | 5.1 | 6.6 | 15.4 |

**Supplementary Table S2:**
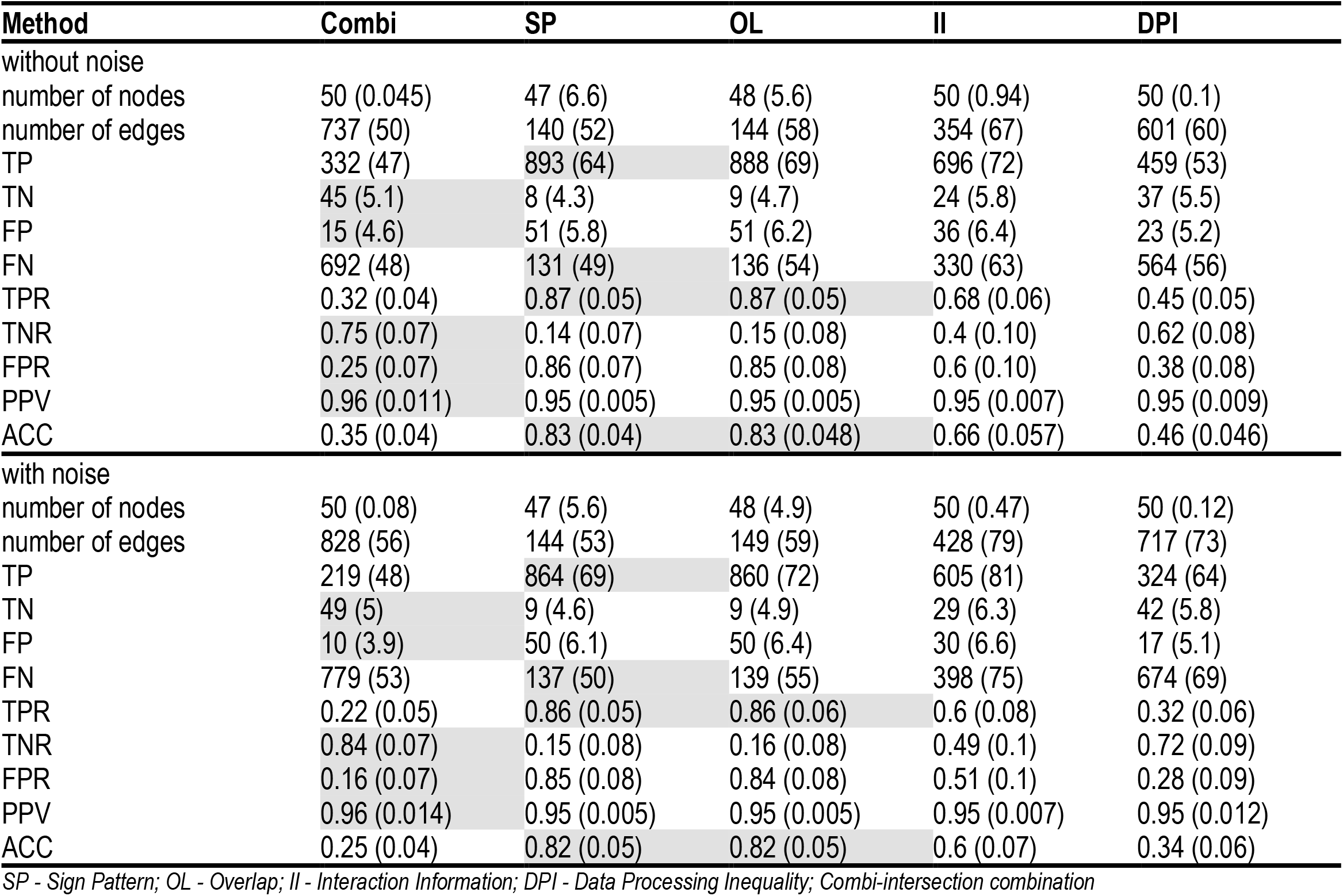
Performance of environmentally-driven edge detection methods on simulated networks. These include 50 microorganisms and 1225 possible associations. Values display median (standard deviation) for simulated networks and simulated networks incorporating noise. Combi refers to intersection combination of all four methods, SP to Sign Pattern, OL to Overlap, II to Interaction Information, and DPI to Data Processing Inequality. The methods with highest (TP, TN, TPR, TNR, PPV, ACC) or lowest (FP, FN, FPR) median, respectively, are highlighted in gray.

**Supplementary Table S3:** Number of triplets a microbial edge is part of in the BBMO network. SP and OL not listed below because they remove 100% of microbial associations that are within at least one triplet. The total number of edges (all) is given along the number of positive (pos) and negative (neg) edges. Combi refers to intersection combination of all four methods, II to Interaction Information, and DPI to Data Processing Inequality.

| Triplets | all | pos (%) | neg (%) | Combi (%) | II (%) | DPI (%) |
| --- | --- | --- | --- | --- | --- | --- |
| 0 | 4 590 | 4 124 (89.8) | 466 (10.2) | NA | NA | NA |
| 1 | 16 193 | 13 369 (82.6) | 2 824 (17.4) | 1 276 (7.9) | 3 851 (23.8) | 4 560 (28.2) |
| 2 | 8 266 | 6 404 (77.5) | 1 862 (22.5) | 1 048 (12.7) | 3 335 (40.3) | 2 585 (31.3) |
| 3 | 667 | 484 (72.6) | 183 (27.4) | 140 (21.0) | 388 (58.2) | 222 (33.3) |
| 4 | 81 | 56 (69.1) | 25 (30.9) | 22 (27.2) | 75 (92.6) | 25 (30.9) |
| 5 | 22 | 20 (90.9) | 2 (9.1) | 2 (9.1) | 22 (100) | 2 (9.1) |
| 6 | 1 | 1 (100) | NA | NA | 1 (100) | NA |

**Supplementary Table S4:** The BBMO network based on real data. The BBMO network contained bacteria (B) and eukaryotes (E) from the picoplankton (p) and nanoplankton (n). This table summarizes the number and fraction of microbial associations classified by EnDED as environmentally-driven. Combi refers to the intersection combination of all four methods, II to Interaction Information, and DPI to Data Processing Inequality. Both methods, Sign Pattern and Overlap, are not shown because both remove all microbial edges found in at least one triplet. For example (last row), 349 (14.9%) associations between bacteria from the picoplankton with eukaryotes from the nanoplankton were classified by intersection combination as environmentally-driven (indirect), II classified 30.6% and DPI 37.2% as environmentally-driven.

| Type | edges | positive | negative | triplets | Combi | II | DPI |
| --- | --- | --- | --- | --- | --- | --- | --- |
| nB | 6 377 | 5 453 (85.5) | 924 (14.5) | 5 150 (80.8) | 376 (5.9) | 1 512 (23.7) | 1 080 (16.9) |
| n+pB | 5 191 | 4 069 (78.4) | 1 122 (21.6) | 4 824 (92.9) | 440 (8.5) | 1 381 (26.6) | 1 678 (32.3) |
| pB | 2 832 | 2 053 (72.5) | 779 (27.5) | 2 160 (76.3) | 125 (4.4) | 569 (20.1) | 631 (22.3) |
| nE | 1 319 | 1 163 (88.2) | 156 (11.8) | 1 016 (77.0) | 113 (8.6) | 350 (26.5) | 254 (19.3) |
| n+pE | 1 165 | 976 (83.8) | 189 (16.2) | 1 006 (86.4) | 158 (13.6) | 353 (30.3) | 370 (31.8) |
| pE | 895 | 820 (91.6) | 75 (8.4) | 543 (60.7) | 44 (4.9) | 153 (17.1) | 113 (12.6) |
| nB+E | 4 703 | 4 080 (86.8) | 623 (13.2) | 4 120 (87.6) | 438 (9.3) | 1 345 (28.6) | 1 043 (22.2) |
| pB+E | 2 520 | 1 908 (75.7) | 612 (24.3) | 1 980 (78.6) | 204 (8.1) | 626 (24.8) | 647 (25.7) |
| nB+pE | 2 483 | 2 100 (84.6) | 383 (15.4) | 2 222 (89.5) | 241 (9.7) | 668 (26.9) | 709 (28.6) |
| pB+nE | 2 335 | 1 836 (78.6) | 499 (21.4) | 2 209 (94.6) | 349 (14.9) | 715 (30.6) | 869 (37.2) |

B - Bacteria; E - Eukaryotes; n - nano fraction; p - pico fraction

